# Conformal Quadrature RF Coils with a Semiopen Architecture for 9.4 T Brain MRI from Mice to Macaques

**DOI:** 10.64898/2026.11.26.769368

**Authors:** Hongyi Yang, Xianghe Ma, Kai Zhong

## Abstract

**Objective:** To develop a semi-open quadrature RF coil architecture that combines brain coverage with low interport coupling at 9.4 T across animal sizes.

**Methods:** Head-conformal high-pass resonators with an adjustable peripheral branch were constructed for four animal head sizes. A shared-rung circuit model, two-capacitance electromagnetic screening, and accepted-power-normalized field analysis guided the implementation. Electromagnetic simulations, loaded bench measurements, saline-phantom comparison with a quadrature birdcage, and ex vivo coronal brain SNR analysis evaluated the macaque implementation.

**Results:** The peripheral branch reduced loaded interport transmission from approximately −10 to −35 dB. Saline-phantom raw SNR medians were 298.9 and 257.8 for the conformal and birdcage coils, respectively (+16.0%). Voxel-normalized pooled ex vivo brain SNR was 648.7 versus 552.8 (+17.3%). Structural imaging demonstrated imaging feasibility in mice, rats/tree shrews, a canine specimen, and macaques, including 176 × 156 × 156 µm^3^ in vivo macaque sampling.

**Conclusion:** The architecture accommodated different animal head sizes through size-specific adjustment while maintaining low interport coupling.

**Significance:** The controlled macaque-scale comparisons support a low-channel-count transceiver option for the demonstrated structural MRI tasks.

## I. Introduction

RF coils determine how efficiently an MRI system excites an anatomical target and detects its signal. The connection between field generation, sensitivity, and noise is established in classical NMR theory^1^. The birdcage resonator provides distributed volume excitation^2^, and quadrature detection combines two appropriately phased transverse sensitivities to improve reception^3^. These principles remain relevant to animal MRI systems with a fixed quadrature interface. For such systems, a practical coil must combine sufficient signal sensitivity, coverage of the intended brain region, and a geometry compatible with animal positioning. Classical birdcage circuit theory relates resonance, excitation, and current patterns^4,5^. Elliptical quadrature designs further show that anatomy-conforming geometry requires an appropriate current distribution and feed arrangement^6^.

At ultra-high field, RF performance depends on both the coil and its dielectric load. Measurements at 3, 7, and 9.4 T have demonstrated that SNR, relaxation times, and transmit-field variation change together with field strength^7^. Although those measurements concern the human brain, they illustrate why an RF design cannot be evaluated from static field strength alone. Animal head dimensions range from the mouse scale to a macaque head approaching the electromagnetic length scale within tissue at approximately 400 MHz. Adapting a coil across these sizes therefore requires reconsideration of conductor geometry, capacitive loading, and feed coupling. Nonhuman-primate studies also require useful whole-brain coverage while preserving access for positioning and physiological support; a quadrature macaque coil developed at 3 T for a stereotaxic head frame exemplifies this requirement^8^. Small-animal RF design must also balance conductor, capacitor, shield, sample, and transmission-line losses^9^. Integrated animal-holder and RF-coil development provides a practical precedent for considering positioning and sensitive volume together^10^.

Open distributed resonators have a substantial design history. The half-birdcage established a half-volume imaging structure^11^, and modified birdcages examined targeted coverage and conductor-current distributions^12^. Birdcage circuit analysis and capacitor-selection methods relate component values to resonance and current patterns^4,5,13^, while a two-dimensional high-pass network demonstrated quadrature operation using inductively coupled resonant elements^14^. Open half-volume TEM coils^15^ and the more recent 7 T opencage^16^ further show that openness and quadrature excitation can coexist. The question addressed here is how a connected high-pass conductor network and a peripheral adjustable branch can be implemented together to obtain useful brain coverage and port isolation. An open birdcage evaluated for mouse cardiac MRI illustrates a related task-specific balance between volume and surface-coil coverage^17^.

Dedicated nonhuman-primate coils offer complementary approaches. *A* quadrature birdcage optimized for squirrel-monkey brain imaging at 9.4 T demonstrates the value of matching volume-coil dimensions to the target^18^. Since the NMR phased-array framework^19^, dedicated monkey receive arrays^20^ and hybrid dipole/loop systems at 10.5 T and 7 T have provided additional sensitivity, acceleration, and RF control^21,22^. *A* recent TBME implementation combines a volume transmitter with an eight-channel marmoset receive array^23^. *A* fixed-quadrature transceiver has fewer independent degrees of freedom and addresses a different hardware requirement. Its performance should be judged against a suitable quadrature reference for the intended task, without implying equivalence to a multichannel array. Flexible and integrated arrays provide further routes to close fitting and system integration^24^, while recent 10.5 T work highlights the importance of coil interfaces and loss control as well as element number^25^.

We developed a family of conformal semi-open high-pass coils using two-port quadrature feeding and an integrated peripheral capacitive branch. The invariant feature is the circuit architecture; geometry and component values are adjusted for each size. The contribution is the joint implementation of semi-open brain coverage and low feed coupling, with the strongest quantitative evidence at the macaque scale. The evaluation proceeds from circuit interpretation and small-animal working-point selection to loaded bench tests, a saline-phantom birdcage comparison, an ex vivo tissue-load comparison, and representative animal-brain imaging. This structure separates measured outcomes from the proposed current-feedback explanation.

## II. Materials and Methods

### A. Coil architecture and size specific implementation

Longitudinal copper strips followed nonmagnetic head-conformal formers and were connected through capacitors at both ends, forming a high-pass distributed resonator (Fig. 1B and Supplementary Fig. S9). Two feeds at symmetric positions on one end were connected to an external 90° hybrid. Each feed was tuned near 400 MHz and matched to the 50 Ω RF interface. An adjustable capacitive branch joined the outer nodes at the opposite end, away from the feed region. Both copper paths between the resonator and the branch capacitor were retained in the physical circuit and in its equivalent representation. Fixed quadrature operation refers to the hybrid-fed two-port resonator, not two independently controlled parallel-transmit channels.

**Fig 1.**
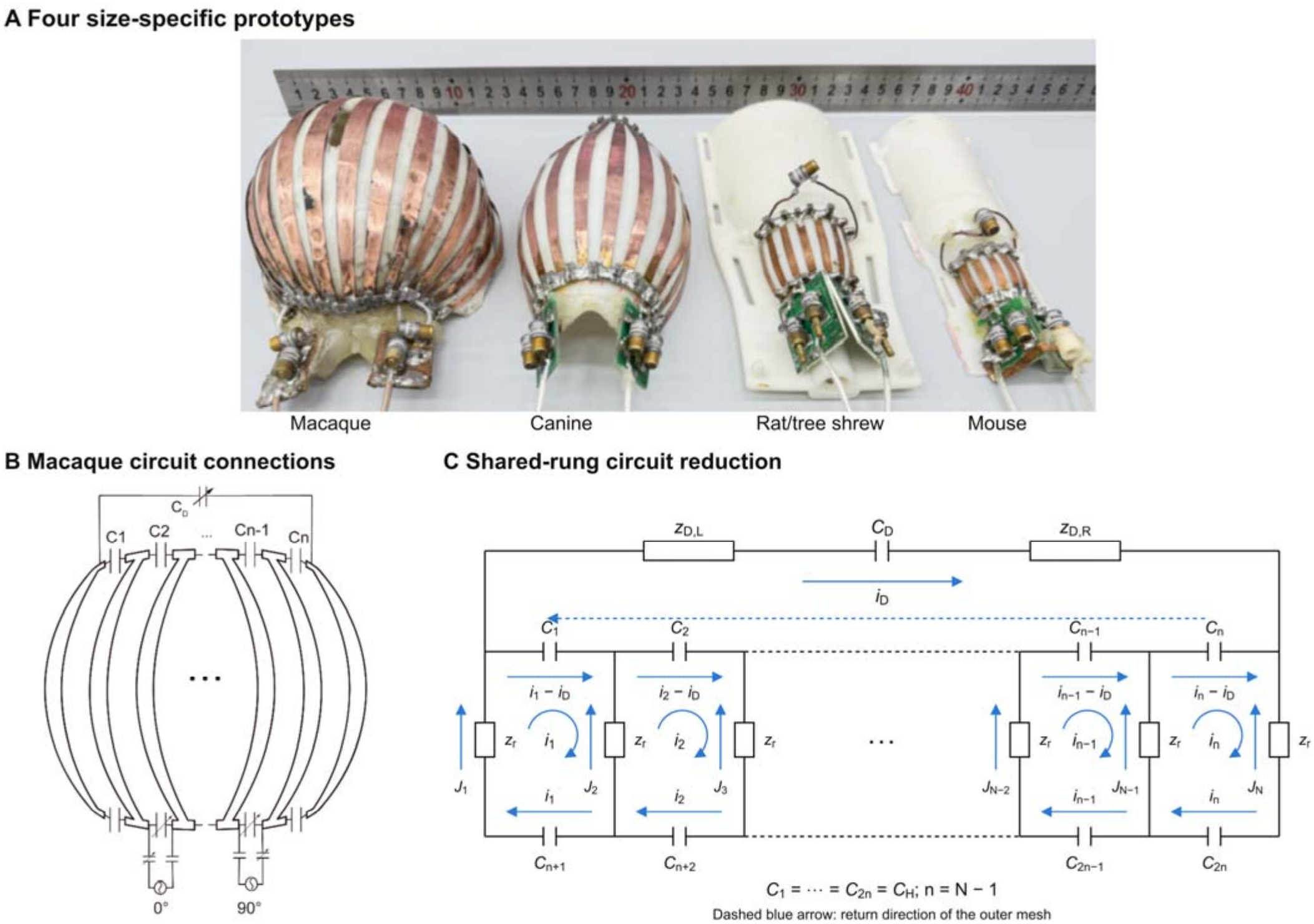
Prototypes and circuit architecture. (*A*) Four constructed sizes, from left to right: macaque, canine, rat/tree-shrew, and mouse. (B) Macaque conductor and feed connections. (C) Unfolded shared-rung model with repeated cells abbreviated by ellipses. The upper capacitors *C*_1,_…,*C*_n_ and lower capacitors *C*_n+1_,…,*C*_2n_ all have capacitance *C*_H_; n = *N* − 1 for *N* rungs. Blue arrows define the current references. The dashed blue arrow shows the outer-mesh return direction along the upper chain, not an additional conductor. Both copper-path impedances flank *C*_D_. Full mouse and macaque layouts are retained in Supplementary Fig. S9.

The mouse baseline used eight conductors, a nominal maximum diameter of 25 mm, approximately 25-mm longitudinal extent, and approximately 3-mm conductor width. The 40-mm, eight-rung implementation supported rat and tree-shrew imaging. *A* 12-rung canine implementation provided an intermediate step before the 120-mm, 16-rung macaque resonator (Table I; Fig. 1A). The same connected high-pass topology was retained, while geometry, end capacitance, matching, and peripheral-branch adjustment were determined for each load. The dimensions and conductor counts describe the constructed size-specific prototypes. Conductor spacing accommodated the target and mounting space; the reported electromagnetic sweep guided capacitance selection rather than an exhaustive optimization of rung number or width. The macaque end capacitance was approximately 6.8 pF.

**TABLE 1.** DESIGN PARAMETERS OF COILS.

| **Implementation** | **Diameter / length (mm)** | **Rungs** | ***C*<sub>H</sub> (pF)** |
|---|---|---|---|
| Mouse | 25 / 25 | 8 | 27 |
| Rat/tree shrew | 40 / 40 | 8 | 20 |
| Canine | 80 / 130 | 12 | 12 |
| Macaque | 120 / 160 | 16 | 6.8 |
| Reference birdcage | 110 / 110 | 16 | 8.2 |

*C*<sub>H</sub> denotes the nominal end capacitance. Geometry and capacitance were selected for each target size; final tuning and decoupling were adjusted under load.

The macaque reference was a quadrature birdcage with an inner diameter and length of 110 mm. The comparison assessed these practical implementations under the same load, rather than isolating a single geometric change. RF shielding was included in the simulation setups shown in the source models; the physical shield and its spacing are relevant because they affect the conductor impedances and fields^26^.

### B. Circuit model and peripheral branch decoupling

The shared-rung circuit in Fig. 1C explains how the peripheral branch enters the high-pass ladder. Let *N* be the number of rungs and n = *N* − 1 the number of cells. The upper end capacitors are numbered *C*_1,_…,*C*_n_ and the lower end capacitors *C*_n+1_,…,*C*_2n_; all have the same capacitance *C*_H_. Each internal mesh current *i*_*k*_ is clockwise. The outer mesh current *i*_D_ flows from left to right through the peripheral branch and returns from right to left along the upper capacitor chain. With the physical rung current *J*_*k*_ directed upward, adjacent meshes give

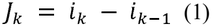

where *i*_0_ = *i*_N_ = 0. For this equal-element model, each rung has self-impedance *z*_r_ = *R* + jωL and each end capacitor has impedance *z*_c_ = 1/(jω*C*_H_). The current through upper capacitor *C*_k,_ referenced from left to right, is *i*_*k*_ − *i*_D,_ whereas the corresponding lower-capacitor current is *i*_*k*_ from right to left. The common −*i*_D_ term follows from the chosen outer-mesh return path; the capacitor currents still differ because the local *i*_*k*_ values differ. Applying Kirchhoff’s voltage law to cell *k* gives

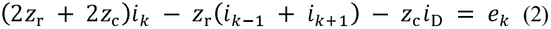

Here *e*_*k*_ represents the cell excitation. The lumped formulation uses identical rung self-impedances; spatial loading and inductive coupling are included in the electromagnetic model. The peripheral path contains both copper sections and CD, represented together by zD. Its voltage must equal the sum of the upper-chain capacitor voltages:

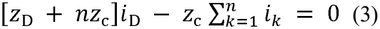

Equation (3) gives *i*_D_ = *z*_c_ Σ *i*_*k*_/(*z*_D_ + n *z*_c_), with the sum over all n cells. Substituting this current into (2) shows how the branch feeds back into the ladder. Let *A* be the mesh-impedance matrix without the peripheral branch: its diagonal entries are 2*z*_r_ + 2*z*_c,_ its adjacent off-diagonal entries are −*z*_r,_ and its other entries are zero.

After eliminating *i*_D,_ the effective matrix has entries

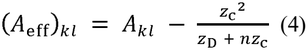

for *k*,l = 1,…,n. Changing *C*_D_ changes *z*_D_ and hence the common feedback term in (4). This alters the cell-current solution and the transfer between the two feeds. The connection to the measured S parameters can be made explicit by retaining the two feed currents and the peripheral-loop current while eliminating the remaining internal currents. Let *Z*_pq,b_ denote the resulting feed-block coefficients, *K*_*p*_ the reciprocal coupling between feed p and the loop, and *Z*_DD_ the loop self-impedance, for p,*q* = 1,2. These coefficients include the fixed feed/matching connections and the rest of the resonator. In particular, *Z*_DD_ includes *C*_D_ and both copper paths; it is not the impedance of *C*_D_ alone. The loop has no independent voltage source, so 0 = *K*_1_ *I*_1_ + *K*_2_ *I*_2_ + *Z*_DD_ *i*_D_. Substituting *i*_D_ = −(*K*_1_ *I*_1_ + *K*_2_ *I*_2_)/*Z*_DD_ into the feed equations gives

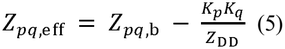

The second term is the branch-mediated contribution to each port impedance. For p ≠ *q* it can partly cancel the direct mutual impedance *Z*_pq,b_; for p = *q* it also changes the input impedance. With the same real reference impedance *Z*_0_ = 50 Ω at both feeds, the standard impedance-to-scattering transformation gives

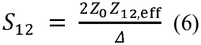

where Δ = (*Z*_11,eff_ + *Z*_0_)(*Z*_22,eff_ + *Z*_0_) − *Z*_12,eff_*Z*_21,eff_. The passive reciprocal circuit has *Z*_12,eff_ = *Z*_21,eff_ and therefore *S*_12_ = *S*_21_ at the same reference planes. *A* small residual mutual impedance gives low interport transmission provided the input match is maintained. Adjustment of *C*_D_ changes the magnitude and phase of the branch contribution, but can shift resonance and matching as well; the working point was therefore selected jointly with tuning and matching. This circuit relation explains how the branch can control *S*_12_. The spatial *B* ^+^ distribution is assessed separately by electromagnetic simulation, rather than inferred from port isolation.

### C. Electromagnetic modeling and working point selection

Three-dimensional frequency-domain electromagnetic simulations were performed near 400 MHz in COMSOL Multiphysics. The explicit model contained the curved copper rungs, distributed end capacitors, peripheral branch, two feeds, support, shield, and homogeneous load; a finite-element mesh is shown in Supplementary Fig. S1. Copper conductivity was 5.8 × 10□ S/m. Unlike the reduced circuit, the field model included the spatial coil–load interaction, which becomes important as electrical size increases^27,28^. The mouse optimization load was an ellipsoid with semi-axes 15 × 9 × 9 mm, relative permittivity 57.1, and conductivity approximately 0.9 S/m.

Distributed end capacitance *C*_H_ and peripheral capacitance *C*_D_ were swept jointly at 400 MHz (Fig. 2). For each combination, target-ROI field amplitude and CV were assessed together with *S*_11_ and *S*_21_. Candidate regions first had to retain continuous target coverage; a low-coupling state and feasible loaded matching were then sought within those regions. The nominal *C*_H_ = 27 pF, *C*_D_ = 3 pF state was selected for the mouse prototype. Field maps at the selected setting, the local-SAR distribution, and a same-scale birdcage comparison are retained in Supplementary Figs. S2 and S3. This workflow selected a usable joint working point rather than the separate extreme of each metric.

The rotating transmit component was defined as 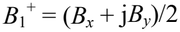 in the specified coordinate/excitation convention, with *B*_*x*_ and *B*_*y*_ transverse to *B*_0_. Field magnitudes were normalized to 1 W total accepted power. For power-normalized incident and outgoing vectors a and b,

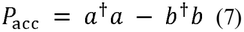

where b = Sa. This normalization includes the simultaneous two-port excitation and does not equate accepted power with sample-only absorption. Transmit-field RMS and CV were calculated within a stated ROI; RMS was divided by the square root of accepted power, and CV was the spatial SD divided by the spatial mean of 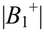. In mouse and macaque birdcage comparisons, the upper-side target ROI near the conformal coil was used for both designs. This was a regional comparison, not a whole-load uniformity test.

The representative mouse/macaque field comparison used brain-equivalent loading and extracted transverse sections, depth profiles, and longitudinal profiles (Fig. 3). The macaque model used relative permittivity 57.1 and conductivity 0.80 S/m. These profiles assessed continuity and rotating-field composition within each model rather than equal field amplitude across different coil sizes. The macaque birdcage comparison instead used an 80-mm saline sphere with relative permittivity 76 and conductivity 1.5 S/m, corresponding to the physical 0.9% saline phantom (Fig. 4). Both coils were assessed in the same target-side upper-half ROI. *A* separate 90-mm brain-equivalent sphere, with density 1000 kg/m^3^, was used for local SAR. For peak complex electric field *E*_pk,_ local SAR was σ|*E*_pk_|^2^/(2ρ), equivalently σ|*E*_rms_|^2^/ρ for an RMS field. Local-SAR maps are supplied in Supplementary Fig. S6; they describe spatial deposition rather than 10-*g*-averaged SAR or sequence-specific temperature rise.

**Fig 3.**
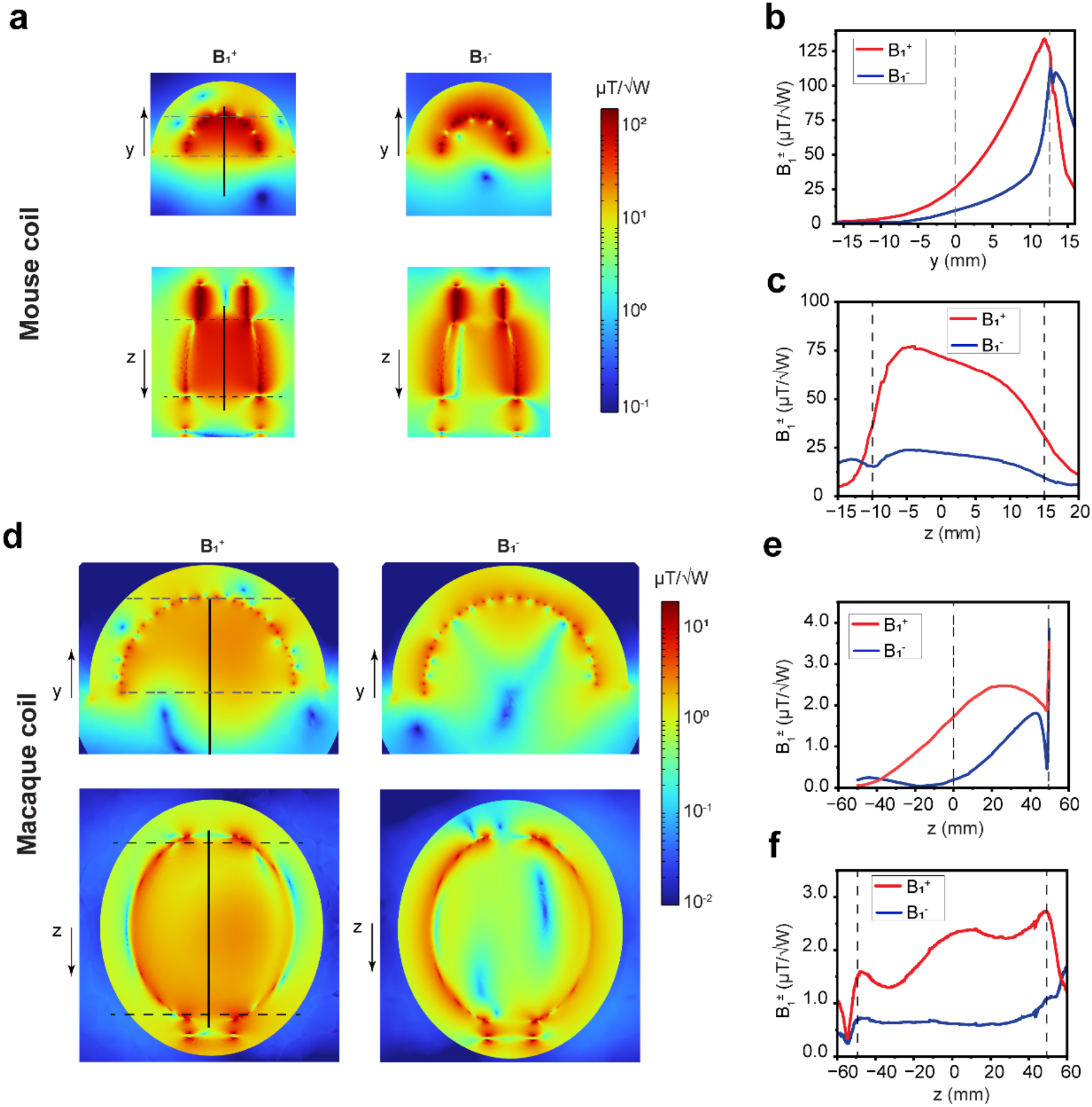
Representative mouse- and macaque-scale electromagnetic fields. (a–c) Mouse and (*d*–f) macaque field sections and depth/longitudinal profiles. Field amplitudes use the 1-W total accepted-power normalization. Dashed lines identify the target extents along the profiles. *B*_1_^−^ is the counter-rotating component under the same transmit excitation. Different spatial coordinates and field scales are retained for the two model sizes.

**Fig 4.**
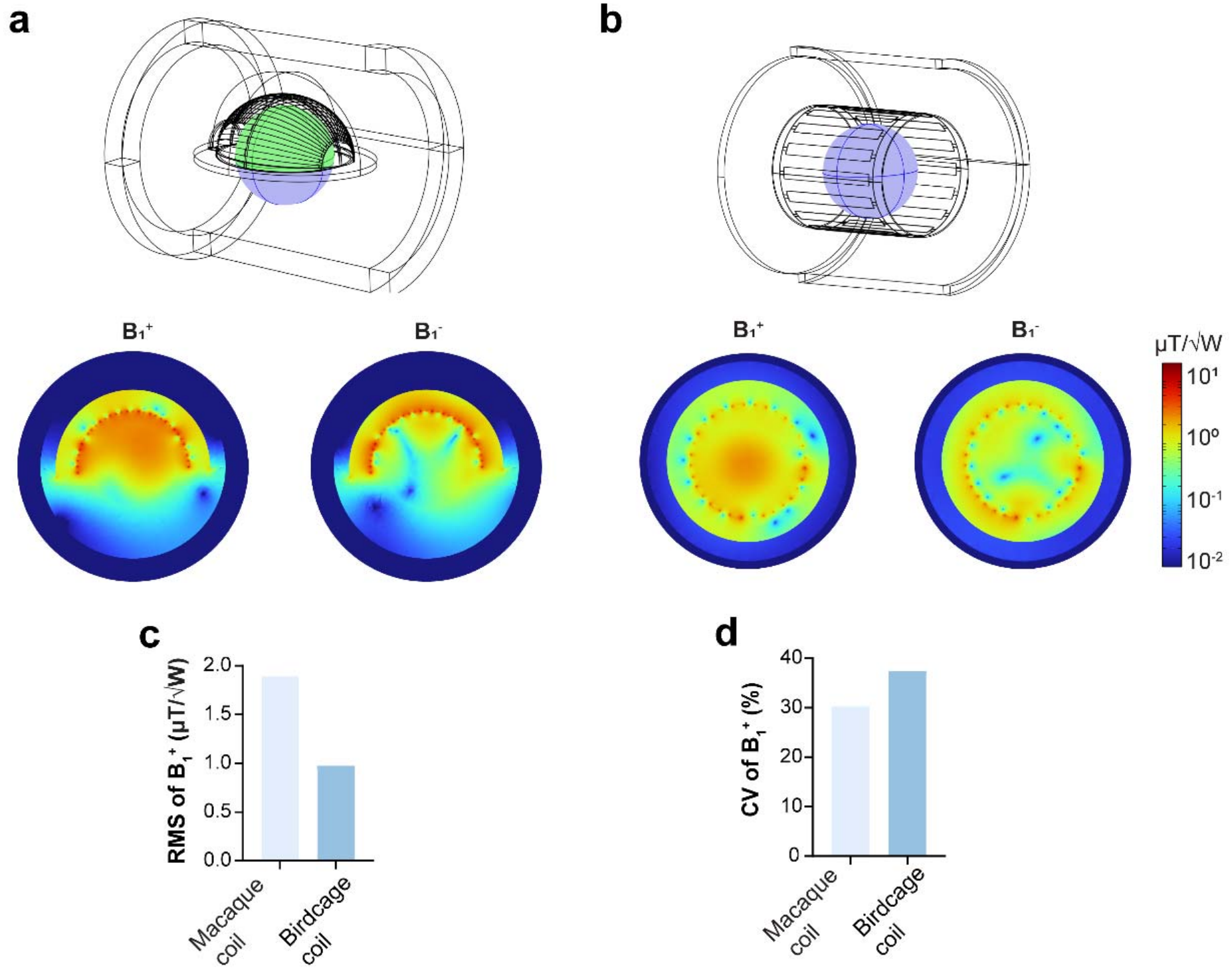
Macaque-scale conformal and birdcage electromagnetic comparison. (a,b) Coil/shield geometry, the shared 80-mm saline load, and corresponding *B*_1_^+^ and transmit *B*_1_^−^ maps. The green upper-half region in (a) defines the target side used for both coils. (c) *B*_1_^+^ RMS and (*d*) CV within this ROI, with equal total accepted-power normalization. The saline parameters are ε_r_ = 76 and σ = 1.5 S/m; this is not the separate brain-equivalent SAR load.

### D. Loaded bench measurements

*A* vector network analyzer measured input reflection and interport transmission near 400 MHz. The mouse implementation was adjusted under saline loading. At the macaque scale, the same approximately 80-mm saline sphere was used for the birdcage, conformal coil without the peripheral branch, and conformal coil with the adjusted branch (Fig. 5). Each configuration was separately tuned and matched. The test consequently assessed the realizable loaded operating states; it was not an otherwise unchanged component-removal experiment. Table II reports the resulting S parameters.

**TABLE 2.** LOADED PORT RESPONSE NEAR 400 MHz.

| **Configuration** | ***S*<sub>11</sub> (dB)** | ***S*<sub>22</sub> (dB)** | ***S*<sub>21</sub> (dB)** |
|---|---|---|---|
| Mouse conformal | −34 | −36 | −47 |
| Macaque birdcage | −21 | −21 | −24 |
| Macaque, no branch | −32 | −32 | −10 |
| Macaque, adjusted branch | −38 | −37 | −35 |

Approximate measured values; macaque states use the same saline sphere and separate loaded adjustments.

**Fig 5.**
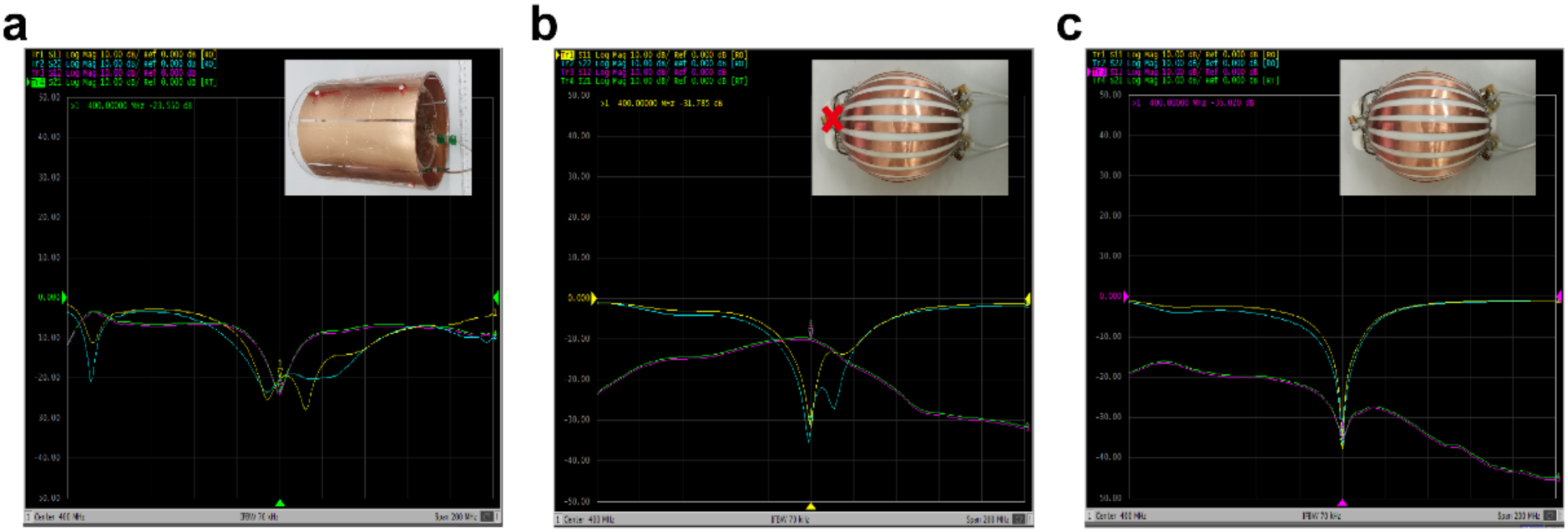
Macaque-scale loaded S-parameter measurements using the same approximately 80-mm saline sphere. Source panels (a), (b), and (c) show the quadrature birdcage, conformal coil without the peripheral branch, and conformal coil with the adjusted branch. Configurations were separately tuned. Table II gives approximate values at 400 MHz.

### E. Saline phantom MRI comparison

MRI was performed on a 9.4 T Bruker BioSpec system. The same saline sphere, acquisition settings, receiver gain, and analysis procedure were used for both macaque-scale coils, with reference RF power calibrated separately. The double-angle method used nominal 45° and 90° FLASH acquisitions with a repetition time sufficiently long for longitudinal recovery^29,30^. The actual flip angle and relative scaling factor β were defined by

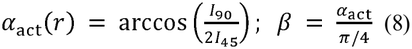

with α_nom_ = π/4 for the mapping pair. Invalid signal ratios were excluded from angle estimation. β describes the calibrated flip angle, whereas the power-normalized simulated 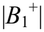 describes transmit efficiency. The same definition was applied to the archived rat- and macaque-brain maps (Fig. 6), which assess calibrated anatomical coverage. These measurements were not treated as spatially matched validation of homogeneous-load simulations. *A* nominal 10° FLASH acquisition and a matched zero-RF acquisition supplied raw image SNR and its noise scale. Because the acquisition and reconstruction affect MRI SNR^31^, a secondary, explicitly defined index was calculated voxelwise as

**Fig 6.**
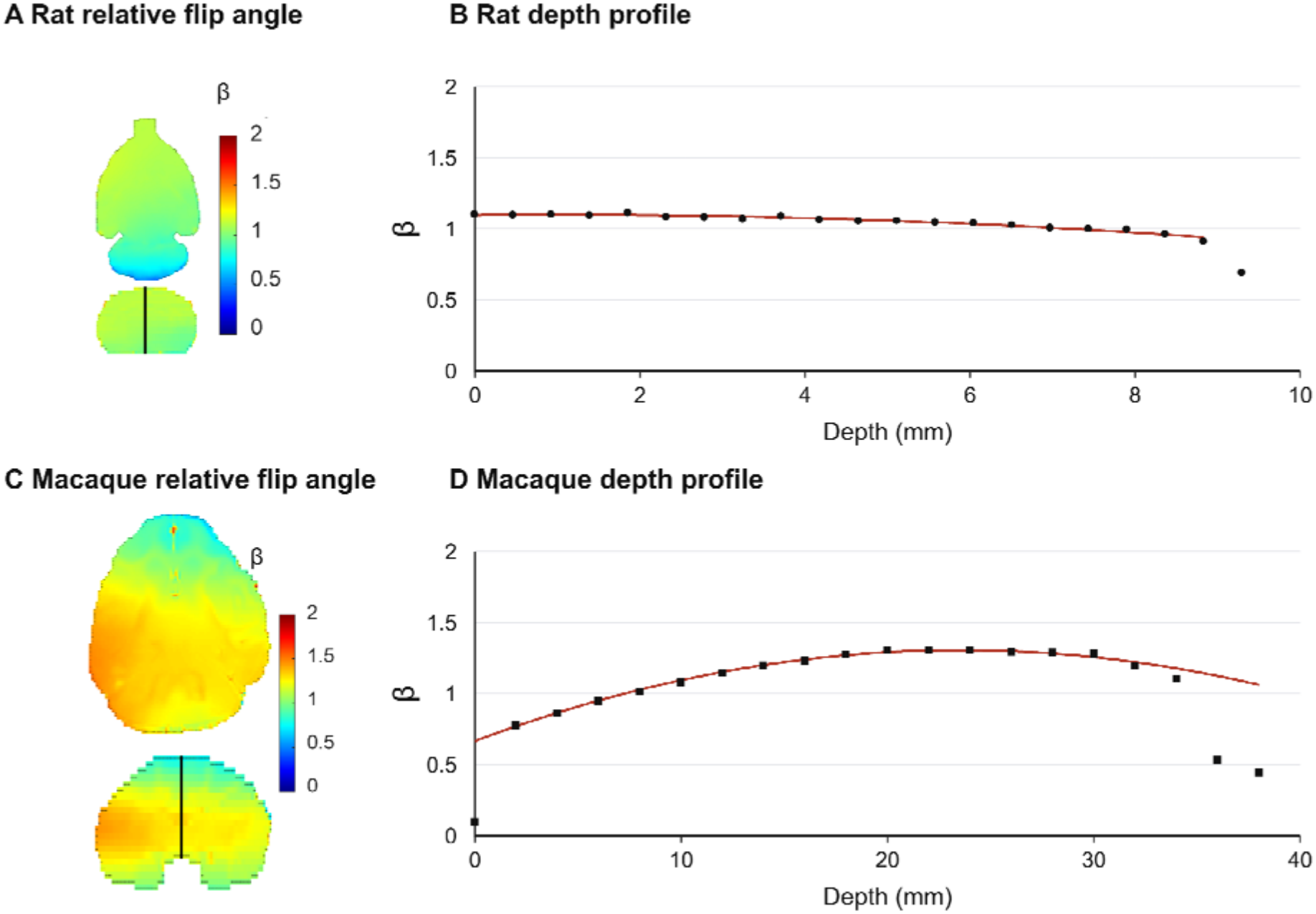
Calibrated relative flip-angle maps and depth profiles in rat (*A*,B) and macaque brain (C,D). β = α_act_/α_nom_. Map color bars and profile axes share a β range of 0–2. Black lines in the lower map sections identify the profile directions. Symbols and red curves retain the archived profile samples and their associated smooth traces. The depth ranges differ with anatomical size. β describes the calibrated excitation state rat er than unit-power transmit efficiency or receive sensitivity.

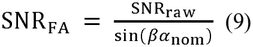

where αnom is 10° in this expression. Voxels with valid signal and 0.2 < β < 1.8 were included. The sine correction reduces the explicit local flip-angle dependence under the intended low-angle conditions, while retaining steady-state and receive-chain contributions. It is therefore termed flip-angle-normalized SNR rather than intrinsic receive SNR. Spatial medians and interquartile ranges describe the effective phantom masks (Fig. 7 and Supplementary Fig. S8).

**Fig 7.**
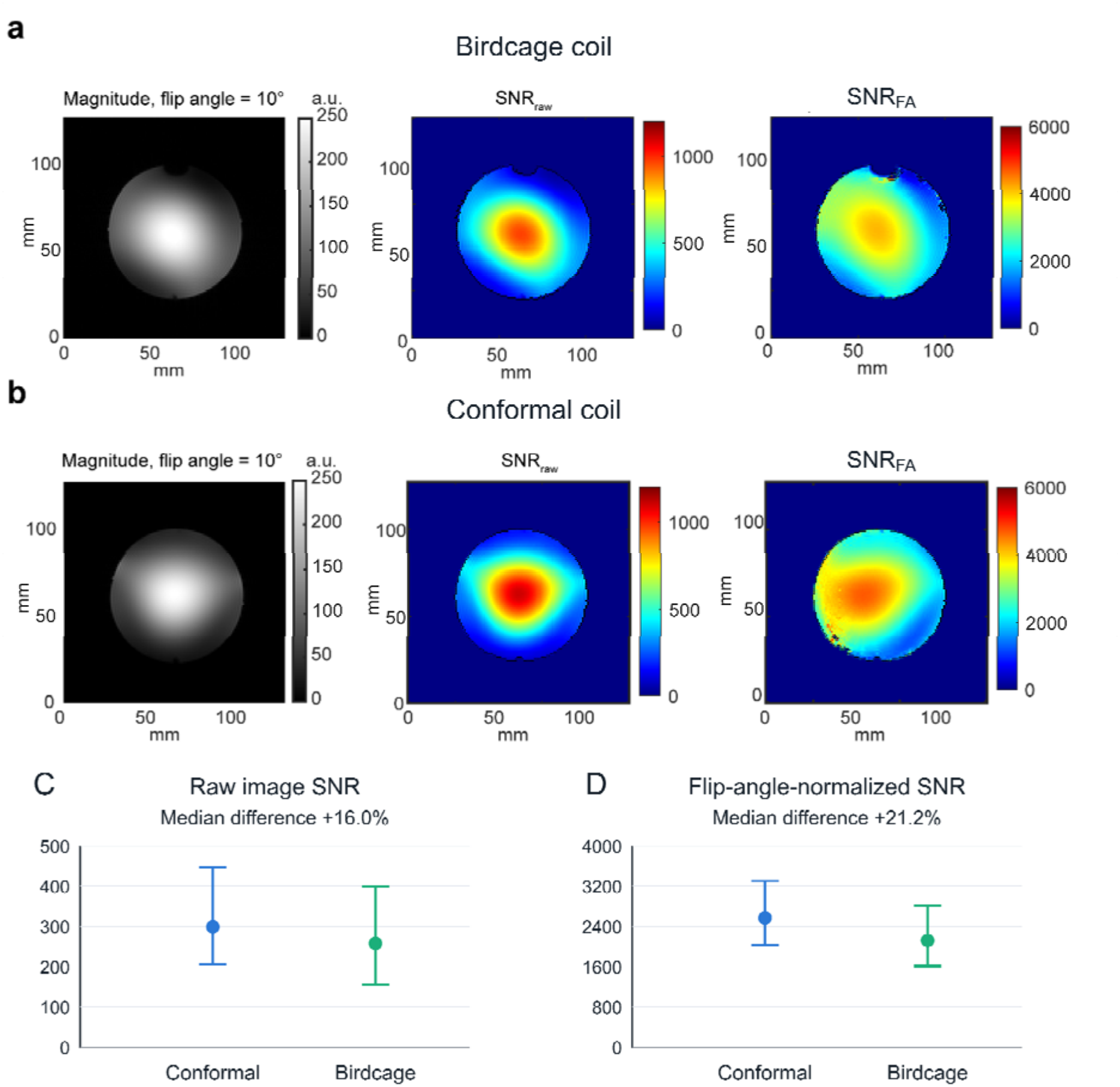
Saline-phantom comparison. (a) Birdcage and (b) conformal: nominal 10° FLASH magnitude, raw SNR, and flip-angle-normalized SNR maps. (C,D) Spatial medians and interquartile ranges, not repeat-scan uncertainty. Valid signal and 0.2 < β < 1.8 define the correction mask. The raw and normalized-SNR medians differ by 16.0% and 21.2%, respectively.

### F. Ex vivo coronal brain comparison

The same ex vivo macaque brain specimen was scanned after changing coils, with preservation and temperature conditions maintained. Coronal FLASH used TR/TE = 500/4 ms, nominal flip angle 30°, a 128 × 128 matrix, 2-mm slice thickness, 100-kHz receiver bandwidth, and one average. FOV was 150 × 150 mm^2^ for the conformal coil and 140 × 140 mm^2^ for the birdcage, giving voxel volumes of 2.7466 and 2.3926 mm^3^. All ten coronal frames per acquisition were assessed on their native reconstruction grids.

After visual review, brain masks used a common threshold of 20% of each frame’s 99.5th-percentile magnitude, with a foreground floor of eight estimated complex-noise SDs. Components smaller than 20 mm^2^ were removed; frames retaining less than 50 mm^2^ were excluded. Masks were neither dilated nor hole-filled. Four 16 × 16-pixel background corner ROIs were placed eight pixels inside the original FOV boundaries, disjoint from brain tissue. All retained brain voxels were pooled for the mean signal, and all corresponding background voxels were pooled for their sample magnitude SD. The reported index was

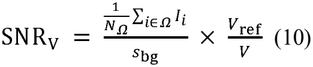

where *N*_Ω_ is the total retained brain-voxel count, *s*_bg_ is the pooled background magnitude SD, and *V*_ref_ is the birdcage voxel volume. Each brain voxel had equal weight; slice means were not averaged with equal slice weights. The volume factors were 0.8711 and 1 for conformal and birdcage acquisitions. This adjustment addresses first-order voxel-volume signal differences and does not turn the acquisition into a pure receive-sensitivity measurement. Thresholds of 25%, 30%, 35%, and 40% supplied sensitivity checks. The image sets were not assumed to be anatomically registered section pairs.

For display only, a shared two-dimensional Fourier-domain notch mask reduced periodic stripes in reconstructed magnitude images, and birdcage panels were rotated in-plane. Display used a common noise/volume-normalized scale (Fig. 8). Mask generation and all numerical SNR calculations used the original unfiltered, unrotated images. The display filter is not a raw complex-*k*-space reconstruction correction.

**Fig 8.**
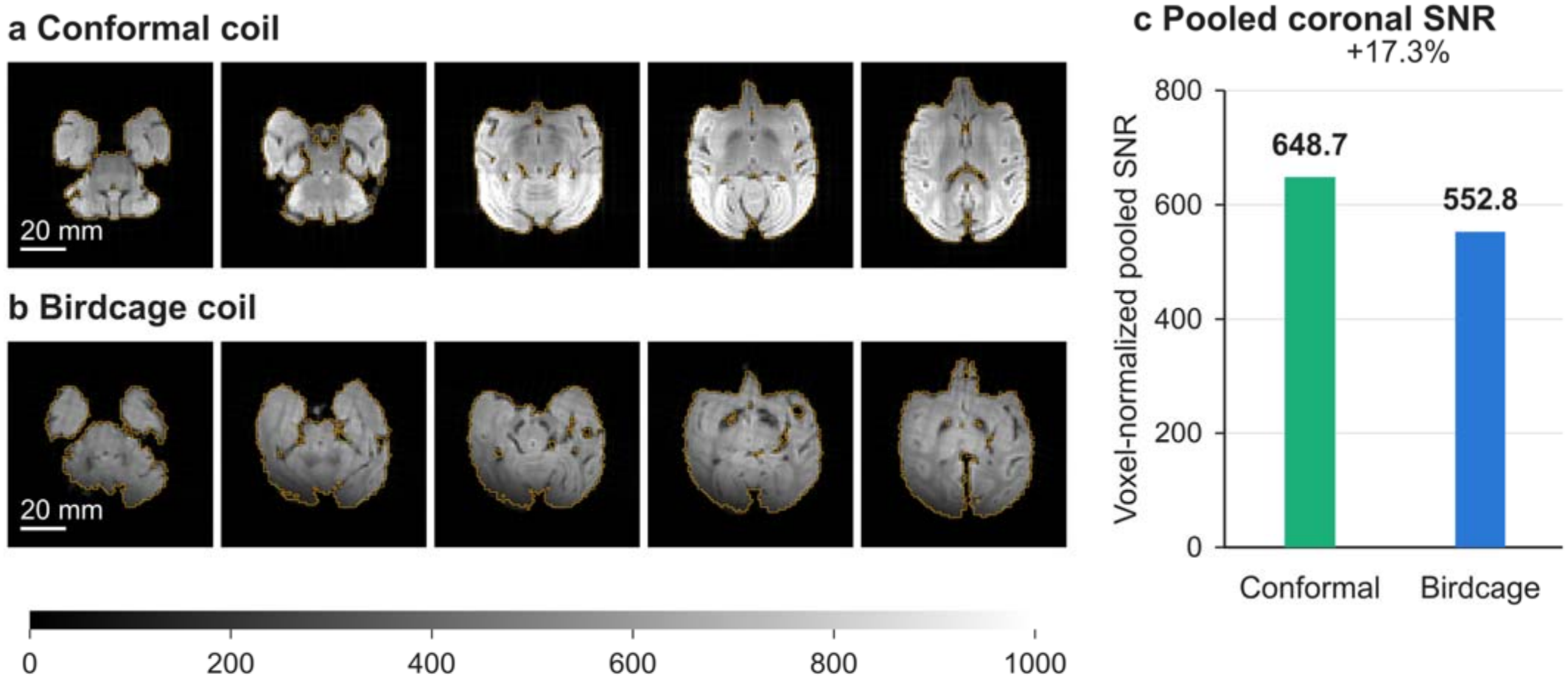
Ex vivo coronal comparison with 20% brain masks. (a) Conformal frames 1,3,5,7,9; (b) birdcage frames 10,9,8,7,6. Yellow contours mark signal masks. Noise ROIs (not shown) are four 16 × 16-pixel original-FOV corner regions per frame, inset 8 pixels and disjoint from tissue. Display copies use a shared Fourier-domain notch filter and noise/volume normalization, with the common 0–1000 scale; birdcage images are rotated. Scale bars: 20 mm. (c) Pooled native-data SNRV over all ten frames: 648.7 and 552.8 (+17.3%). Statistics use unfiltered, unrotated data; are not anatomically registered pairs. Bars summarize one acquisition per coil.

### G. Animal brain imaging and reporting

Representative structural images documented the four size-specific implementations (Fig. 9); rat imaging and the mouse phantom/brain validation are also retained in Supplementary Fig. S12. Mouse-coil phantom GRE used TR/TE = 1000/7.3 ms, nominal flip angle 60°, 0.1 × 0.1-mm^2^ in-plane sampling, and 1-mm thickness. The corresponding thesis mouse-brain GRE used TR/TE = 2500/6.2 ms and nominal flip angle 75°. Tree-shrew three-dimensional FLASH provided approximately 75-µm isotropic sampling. Archived ex vivo canine FLASH used TR/TE = 100/16 ms, flip angle 18°, FOV 90 × 60 × 54 mm^3^, and matrix 750 × 500 × 450, giving 120-µm isotropic sampling. Different specimens and protocols illustrate coverage at successive sizes; their signal levels were not compared as a cross-species performance ranking.

**Fig 9.**
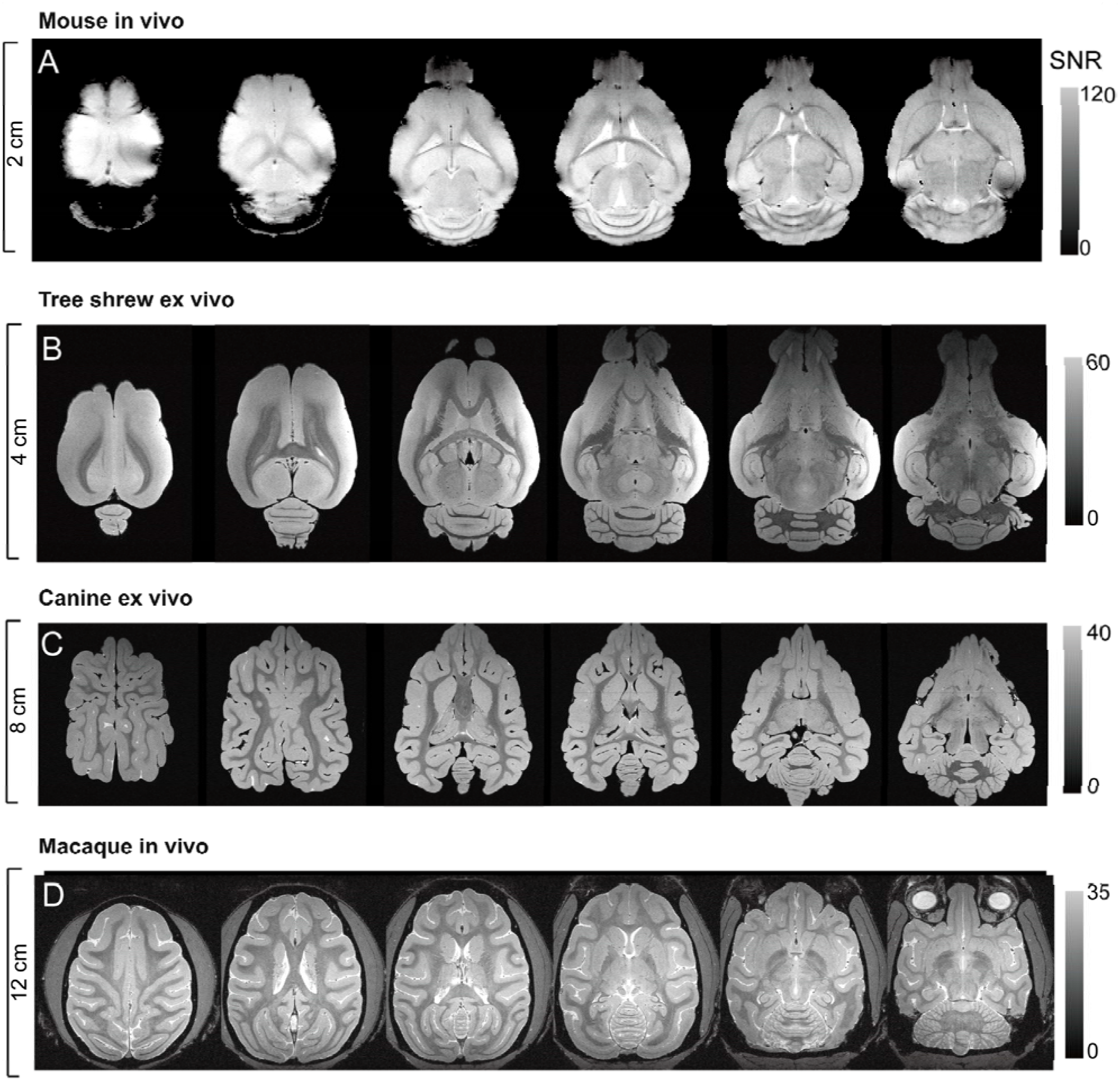
Representative brain MRI across four coil sizes. (*A*) Mouse in vivo, (B) tree shrew ex vivo, (C) canine ex vivo, and (D) macaque in vivo. Spatial brackets and gray bars retain the labels of the archived montage, with image-SNR display ranges of 0–120, 0–60, 0–40, and 0–35, respectively. These examples document structural coverage under different specimens and protocols; the controlled between-coil SNR comparisons are shown separately in Figs. 7 and 8.

In vivo macaque three-dimensional structural imaging used FOV 90 × 80 × 80 mm^3^ and matrix 512^3^, giving approximately 176 × 156 × 156 µm^3^ sampling (Fig. 10). Ex vivo three-dimensional images and local enlargements assessed ventricular, callosal, and lateral-geniculate-region detail. Layer-like signal patterns were described without histological assignment. Quantitative coil comparisons were descriptive: phantom voxels and sections of one specimen were not treated as independent biological replicates, and no coil-comparison P values were constructed from them.

**Fig 10.**
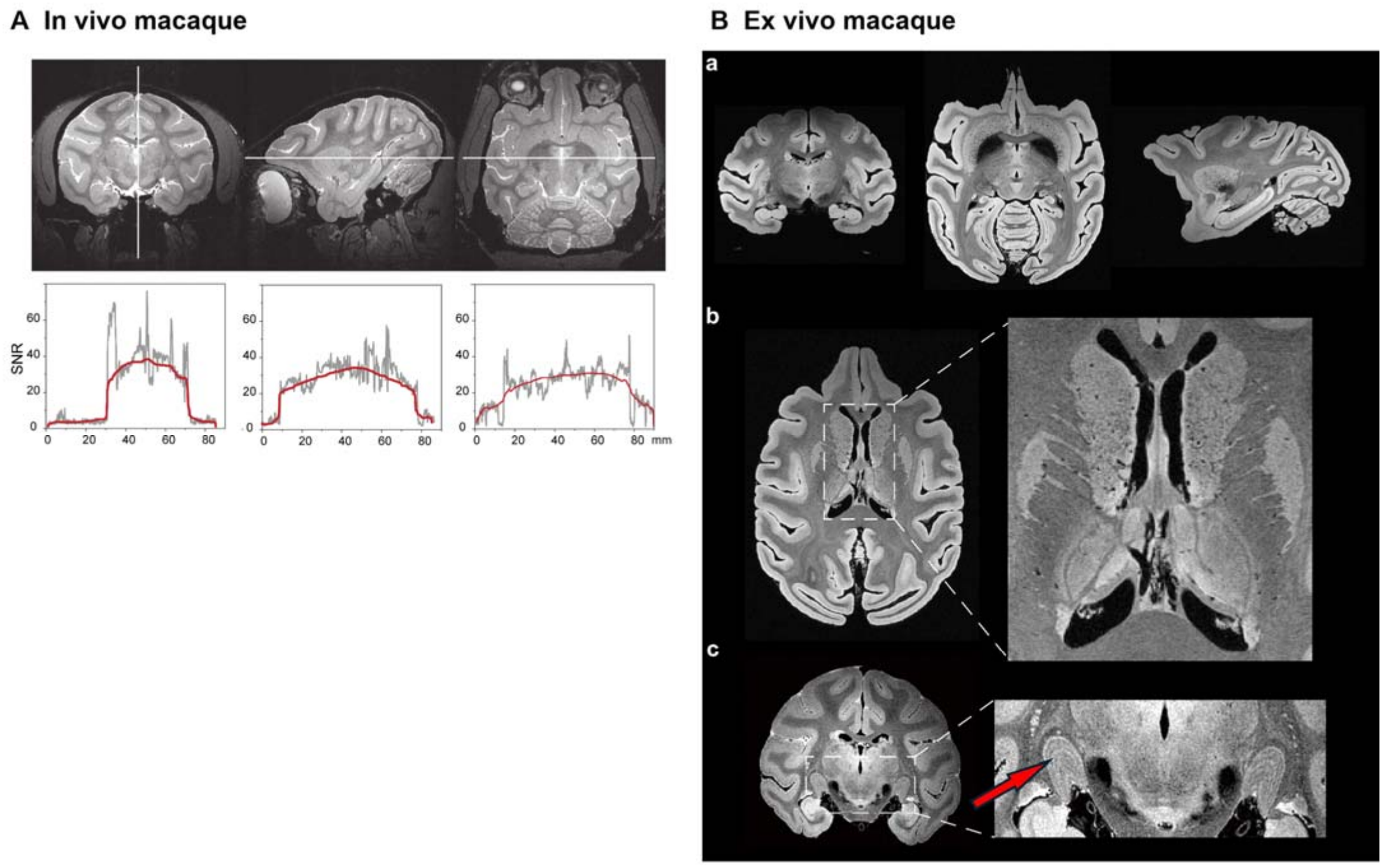
Macaque structural imaging with the conformal coil. (*A*) In vivo orthogonal reconstructions at approximately 176 × 156 × 156 µm^3^; source profiles show apparent image SNR along marked lines (gray: original, red: smoothed spline). (B) Ex vivo reconstructions and ventricular/callosal/lateral-geniculate-region enlargements. Arrows mark layer-like contrast, without histological layer assignment. The in vivo and ex vivo acquisitions demonstrate different imaging conditions.

## III. Results

### A. Working point and size specific implementation

The two-capacitance sweep showed distinct amplitude, CV, and port-response optima rather than a single common optimum (Fig. 2). Changing *C*_H_ modified the principal resonant paths, while *C*_D_ also shifted the field pattern and interport response. The selected approximately 27-pF/3-pF region retained useful target-region amplitude and spatial continuity with adjustable port coupling. At this state, the mouse field maps showed coverage through the target load, and the same-scale comparison retained an efficiency– uniformity tradeoff: the conformal design had higher target-ROI field RMS, whereas the reference birdcage had lower CV (Supplementary Figs. S2 and S3). The fabricated mouse coil reached approximately −34/−36 dB input reflection and −47 dB *S*_21_ under the documented load.

At the representative mouse and macaque sizes, the simulated 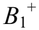 distributions remained continuous through the selected target regions, with greater spatial variation in the larger model (Fig. 3). The depth and longitudinal profiles therefore support size-specific adjustment of the shared architecture, not geometrically proportional scaling with unchanged performance. 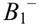 in these simulations is the counter-rotating component of the same applied excitation; its lower amplitude over much of the target region complements the assessment of the chosen quadrature drive.

In the macaque saline model, the conformal coil concentrated its higher field toward the covered side, whereas the birdcage distribution was more centered (Fig. 4). Within the predefined upper-half ROI, the conformal implementation had higher unit-accepted-power 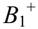 RMS and lower CV than this birdcage. The model geometry and ROI limit this to a regional transmit-field comparison. Together, the sweep and the loaded comparison provide a field-based rationale for the selected design independently of the image-SNR results.

### B. Macaque loaded isolation and field coverage

All three macaque bench configurations achieved input reflection below −20 dB (Fig. 5; Table II). Conformal-coil *S*_21_ was approximately −10 dB without the peripheral branch and −35 dB with it, an approximately 25-dB reduction between the separately tuned configurations. The adjusted coil had *S*_11_/*S*_22_ of approximately −38/−37 dB, compared with −21/−21 dB for the birdcage; birdcage *S*_21_ was approximately −24 dB. These measurements establish the low-coupling working state used for MRI. For the reciprocal two-port coil, this *S*_21_ reduction also represents the corresponding reduction in *S*_12_ at the same reference planes; the measured input reflections distinguish the low-coupling state from attenuation caused by severe mismatch.

The loaded port measurements and the field comparison address complementary parts of the design. The adjusted branch provided the low-transmission working state, while Fig. 4 established the target-region field distribution at the chosen operating point. In the separate brain-equivalent sphere, relatively high local SAR lay near the conductor-side boundary, with lower deposition deeper in the load (Supplementary Fig. S6).

### C. Calibrated brain flip angle maps

The rat and macaque DAM maps showed continuous calibrated flip-angle coverage within the displayed brain sections (Fig. 6). The rat map varied more gradually over its smaller target, while the macaque map exhibited a broader spatial variation. The depth profiles along the marked directions illustrate this spatial variation. These measurements complement the modeled field coverage under anatomical loading. Since the subjects, planes, and reference-power calibrations differ from the homogeneous simulations, they are interpreted as coverage demonstrations rather than a quantitative model–measurement error analysis.

### D. Controlled saline phantom comparison

The two coils had similar calibrated β medians: 0.722 for conformal and 0.707 for birdcage (Supplementary Fig. S8). The β distributions retained different spatial patterns and do not by themselves establish better measured flip-angle uniformity. Raw SNR medians were 298.9 and 257.8, respectively, a 16.0% difference (Fig. 7; Table III). Corresponding interquartile ranges were 206.7–446.5 and 155.9–399.4. Means were 356.0 and 303.0, indicating the same direction of difference. The higher-signal region of the conformal implementation lay toward its covered side rather than the phantom center.

After the one-factor flip-angle correction, SNR_FA_ medians were 2571.1 and 2121.6, a 21.2% difference; respective interquartile ranges were 2018.9–3303.1 and 1612.5–2811.1. CVs of the corrected index were 32.3% and 37.3%. Thus, the median SNR advantage in the evaluated masks remained after reducing the explicit excitation-angle contribution. The correction did not convert this comparison into an independent measurement of intrinsic receive performance.

**TABLE 3.** MACAQUE-SCALE IMAGE-SNR COMPARISON.

| **Metric** | **Conformal** | **Birdcage** | **Difference** |
|---|---|---|---|
| Saline raw SNR median | 298.9 | 257.8 | +16.0% |
| Saline *β* median | 0.722 | 0.707 | +2.1% |
| Saline SNR<sub>FA</sub> median | 2571.1 | 2121.6 | +21.2% |
| Brain SNR<sub>V</sub>, 20% | 648.7 | 552.8 | +17.3% |

Spatial phantom summaries and one tissue acquisition per coil; different indices are not combined. SNR<sub>FA</sub> uses the sine correction, not a full intrinsic-SNR measurement.

### E. Ex vivo tissue load comparison

All ten frames per coil satisfied the inclusion criterion at the 20% threshold. Brain masks retained 17,743 conformal-coil and 17,710 birdcage voxels. The voxel-normalized pooled image-SNR indices were 648.7 and 552.8, respectively, corresponding to a 17.3% higher value for the conformal coil (Fig. 8; Table III). Across the 25%–40% mask thresholds, the relative difference remained positive at 15.8%–16.9%. These are sensitivity analyses of the same acquisitions, not repeated specimens. The tissue-load result complements the saline comparison using a different SNR definition and acquisition; their percentage differences were not combined.

### F. Brain imaging from mice to macaques

The four-size series showed visible brain anatomy from mouse to macaque, including the canine prototype that bridges the intermediate and macaque sizes (Fig. 9). Rat FLASH/RARE and approximately 75-µm-isotropic tree-shrew imaging provide additional intermediate-scale examples (Supplementary Fig. S12). Ex vivo canine images at 120-µm isotropic sampling depict cortical and cerebellar anatomy. The macaque coil provided in vivo whole-brain reconstructions at approximately 176 × 156 × 156 µm^3^, including gray/white-matter boundaries, ventricles, and deep structures (Fig. 10A). Ex vivo macaque images retained local detail in ventricular, callosal, and lateral-geniculate regions (Fig. 10B), described as layer-like contrast without histological assignment.

## IV. Discussion

The study links a shared semi-open high-pass architecture to size-specific electromagnetic design and experimental evaluation. Mouse-scale screening established how field coverage, CV, and port coupling constrain the capacitance choice; representative field simulations then examined the larger macaque load. The four prototypes and brain-image series demonstrate realizations across sizes, including the canine intermediate. The strongest quantitative comparison remains at the macaque scale, where the same-load birdcage experiments showed higher image-SNR summaries for the conformal implementation. These results support the demonstrated configurations without assuming that all sizes share the same homogeneity, efficiency, or SNR gain.

The relationship to prior resonators is important for defining the contribution. Half-birdcage and U-shaped birdcage studies established open targeted coverage^11,12^, while the half-volume TEM used a different distributed resonator for quadrature excitation^15^. The two-dimensional high-pass ladder relied on inductively coupled resonant elements^14^, whereas the present structure has physically shared rungs and capacitor-connected end paths. The opencage designed nonperiodic birdcage cells and currents while retaining a quadrature volume-like structure^16^. Here, the practical design emphasis is a head-conformal half-open conductor network, hybrid-fed symmetric feed locations, and an adjustable connected branch outside the principal feed region. The achieved isolation and macaque-load comparisons distinguish the evaluated implementation; they do not establish that openness, quadrature, or conformality was first introduced here. Supplementary Table S2 summarizes these distinctions without cross-study SNR rankings.

The peripheral branch provides a current path whose contribution can oppose transfer between the feeds. Equation (5) separates the feed-block mutual impedance from the branch-mediated term, and (6) links their residual directly to *S*_12_. Related work has used auxiliary resonators or element-current control to reduce mutual coupling^32,33,34^; the present branch is instead connected to the shared-rung network. The approximately −25dB transmission difference between the separately tuned macaque configurations supports its practical decoupling role. The circuit model gives a plausible explanation for this control, while the loaded field calculations show the coverage retained at the selected setting. Resolving the individual conductor-current contributions would require a loaded current analysis; low *S*_12_ alone does not establish field uniformity or reconstruction of a missing half-birdcage.

Geometry also affects the field and SNR comparison. Bringing conductors closer to the intended region can change both excitation and receive weighting, but filling factor alone does not explain all observed differences. Load permittivity, losses, shield coupling, return paths, and matching contribute together. The simulated upper-side ROI and effective experimental phantom masks specify where the reported advantages were evaluated. Their outcomes should not be generalized to whole-load uniformity. Quadrature performance also concerns two transverse field components relative to *B*_0_, not only a 90° phase setting on the hybrid. Thus, the useful loaded field patterns and measured port state are more relevant than assuming that geometric symmetry guarantees a particular field mode. The simulated and measured field plots therefore serve different roles: homogeneous models support controlled parameter and load comparisons, whereas the rat/macaque β maps show calibrated anatomical coverage. Agreement in broad coverage is informative, but is not a pointwise validation across unlike planes or loads.

The two SNR comparisons answer related but different questions. The saline experiment controls the load and acquisition settings and provides a secondary excitation-angle correction. The ex vivo comparison measures actual tissue-loaded image SNR with pooled brain voxels, background noise, and voxel-volume adjustment. Both describe image performance rather than separately measured receiver sensitivity. The correction in (9) retains FLASH steady-state terms, while the tissue result also depends on spatial coverage and excitation. The common 20% mask was selected by visual tissue-coverage review; persistence of the difference over alternative thresholds provides an internal check against weak-tissue exclusion. The observed advantage applies to these pooled coronal acquisitions of one specimen. Anatomically matched repeat scans would establish its consistency across specimens.

RF power deposition deserves separate interpretation. Power-normalized 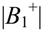 and local-SAR distributions provide design information, but are not interchangeable with SAR efficiency based on a specified mass average. The present homogeneous models and retained local maps support a limited deposition assessment. They do not provide the paired heterogeneous-head SAR_10g_ analysis needed for a direct SAR-efficiency comparison. Consequently, transmit claims are restricted to the defined field/coverage metrics, and image-SNR gains are not presented as improved SAR efficiency. More detailed loads and sequence-average power evaluation would extend this assessment without changing the measured isolation and image findings.

Multichannel receive and parallel-transmit systems remain important routes to acceleration, independent sensitivity weighting, and field control^20,21,22,23^. The present two-port resonator instead supplies a compact fixed-quadrature transceiver for the demonstrated structural tasks. No matched array or birdcage-transmit/separate-receive experiment was available, so neither equivalence nor superiority to those systems is claimed. The macaque result is valuable within that low-channel-count setting: the same broad circuit concept used at smaller sizes was re-tuned for a larger load, obtained low interport transmission, and retained useful brain imaging coverage. Future work can use the structural model to test conductor-current predictions and examine repeatability with anatomically matched acquisitions.

## V. Conclusion

*A* head-conformal semi-open high-pass architecture with an adjustable peripheral branch was implemented for brain MRI from mice to macaques at 9.4 T. The branch provided a low-coupling loaded quadrature working state, while size-specific adjustment maintained useful target coverage. Macaque saline and ex vivo coronal comparisons showed higher image-SNR summaries than the tested quadrature birdcage under the stated conditions. These findings support the architecture as a low-channel-count transceiver option for the demonstrated animal-brain structural imaging tasks.

